# Dysregulated lymphatic remodeling promotes immunopathology during non-healing cutaneous leishmaniasis

**DOI:** 10.1101/2025.05.01.651666

**Authors:** Lucy Fry, Flavia Neto de Jesus, Matheus Batista Carneiro, Hayden Roys, Anne Bowlin, Nathan C Peters, Pierre-Yves von der Weid, Tiffany Weinkopff

## Abstract

Cutaneous leishmaniasis (CL) is a vector borne disease that is endemic to tropical and sub-tropical regions of the world disproportionately affecting those of low socioeconomic status. The combined role of the parasite and the host’s immune response in determining disease severity has made it challenging to discover new anti-leishmanial treatments. Previous work from our lab has established that the dermal lymphatic network is necessary for wound resolution in a model of healing CL with *Leishmania major* parasites. In CL, lymphatic remodeling allows for accumulated fluid to drain from the lesional site, thereby reducing disease severity. In this report, we present a new mechanism of immunopathology during non-healing CL brought about by *L. amazonensis* infection. We show non-healing CL develops alongside an accumulation of cells and fluid in the skin, resulting in chronic inflammation. Lymphatic remodeling is attenuated during the chronic phase of *L. amazonensis* infection. Moreover, the percentage of proliferating lymphatic endothelial cells (LECs) decreases from 6 to 12 weeks post infection (wpi), leading to a decrease over time in lymphatic vessel (LV) density. To induce lymphangiogenesis, exogenous vascular endothelial growth factor-C (VEGF-C) was administered by adenoviral delivery. VEGF-C increased LV dilation leading to reduced lesion sizes without altering parasite burdens, arguing targeting the lymphatics can alleviate immunopathology. Taken together, these results show impaired lymphatic function contributes to non-healing disease due to *L. amazonensis* infection and the lymphatics can be targeted to decrease inflammation in the skin during infection.

## Introduction

Cutaneous leishmaniasis (CL) is the spectrum of diseases caused by vector-transmitted *Leishmania* protozoan parasites endemic to the tropical and subtropical regions of the world. Each year there are 1-2 million new cases of CL with 12 million ongoing infections [1,2]. Based on the species of infecting parasite, different clinical manifestations present within the spectrum of CL, ranging from spontaneously self-healing skin lesions to non-healing permanent disfigurations. Topical, oral, and intravenous medications can manage CL and alleviate symptoms, but since they rarely achieve a sterile cure, many patients remain refractory to treatment [3,4]. Furthermore, there are no human vaccines for CL, emphasizing the need for new therapeutics [5]. Although CL is endemic in tropical regions, rising autochthonous transmission in the US and climate change challenges are predicted to increase CL in North America further highlighting the urgent need for novel therapeutics [6–8].

The species of parasite and the host immune response dictates clinical disease where both overactive and insufficient immune responses can lead to chronic disease [6]. Specifically, control of the intracellular infection is dependent on an effective CD4^+^ Th1 immune response [7]. Th1 cells mediate the production of IFNγ leading to macrophage activation, ultimately resulting in parasite control [8,9]. Inability to initiate a dominant Th1 response is a major cause of chronic non-healing disease where mice developing a Th2 response are more susceptible to *Leishmania* parasites [10]. Moreover, patients presenting with non-healing skin lesions exhibit a mixed Th1/Th2 response [10–12]. For example, during spontaneous resolution of CL caused by *L. major,* which is prevalent through Africa and the Middle East, a predominant Th1 immune response controls parasite burdens thereby inducing lesion resolution [13–15]. In contrast, non-healing and diffuse CL caused by *L. amazonensis*, which is prevalent throughout the Central and South America, leads to a mixed Th1/Th2 immune response with impaired parasite control and persistent lesions [10,11,15,16].

The immune response not only influences the host’s ability to control the parasite but also plays a crucial role in the progression of tissue damage in CL. While the Th1/Th2 balance can determine disease outcome, an excessive inflammatory response can lead to tissue destruction, even in the presence of low parasite burdens [17–21]. However, the precise mechanisms by which the host response contributes to non-healing lesions in CL remain unclear. Given that host-mediated immunopathology underpins the pathogenesis of a variety of inflammatory conditions, many diseases therapeutically target the host response in cancers and autoimmune diseases [22,23].

The lymphatic vasculature regulates the immune response to infection by transporting soluble antigen and antigen presenting cells to the draining lymph nodes (dLNs) and actively modulating inflammatory immune cell exit from the site of inflammation [24,25]. Additionally, immune cells, specifically macrophages, are found in close proximity to lymphatic endothelial cells (LECs) within the skin allowing for the regulation of new lymphatic growth through receptor mediated interactions [24]. During infection with *Leishmania* parasites, inflammatory cells migrate into the skin and aid in the destruction of parasites, tissue repair, and wound healing [25,26]. After the inflammatory insult is resolved, leukocytes exit from the inflammatory site through lymphatic vessels (LVs) to the dLN by responding to chemokines released from LECs [29,30]. Because *L. amazonensis* infection leads to non-healing chronic lesions, we hypothesize these lesions accumulate fluid and immune cells over time and that lesions are unable to drain due to impaired lymphatic expansion and function.

Lymphangiogenesis, or the creation of new LVs, is an essential mechanism for lesion resolution during murine *L. major* infection [27]. We found vascular endothelial growth factor-A (VEGF-A) and vascular endothelial growth factor receptor-2 (VEGFR-2) signaling is the main pathway driving lymphangiogenesis following *L. major* infection with no role for VEGFR-3 signaling [27]. In the *L. major* healing model of CL, macrophages produce VEGF-A to promote lymphangiogenesis, but the role of the lymphatic vasculature in chronic non-healing CL is not known [28].

Because we hypothesize the accumulation of fluid and cells is due to attenuated lymphangiogenesis in non-healing CL, we first characterized the vascular and immune responses using an experimental model of non-healing CL caused by *L. amazonensis* [29,30]. Progressive disease was associated with decreased lymphangiogenesis and increased accumulation of fluid and neutrophils at the site of infection, suggesting lymphangiogenesis is impaired during *L. amazonensis* infection. We sought to adenovirally-deliver VEGF-C to induce lymphangiogenesis following *L. amazonensis* infection. Importantly, adenovirally delivered VEGF-C reduced lesion sizes without altering parasite burdens. This identifies the lymphatics as a new mechanism restricting immunopathology for CL and offers a promising potential target to induce healing in patients suffering from CL, and other inflammatory diseases.

## Results

To investigate mechanisms of immunopathology during non-healing CL, the well-established model of non-healing CL was used. C57BL/6 mice were infected with 100,000 *L. amazonensis* parasites and consistent with other reports, *L. amazonensis* infection resulted in dermal lesions that continually increased in size until the experimental endpoint at 12 weeks post infection (wpi) (Figure 1A) [31]. Lesions measured ∼1.5 mm thick at 12 wpi and presented as elongated inflexible nodules (Figure 1A-B). Consistent with larger lesions, parasite burdens were elevated at 12 wpi compared to 6 wpi (Figure 1C). Increased parasite burdens as well as progressive lesion development suggests the immune response during *L. amazonensis* infection is not sufficient to control parasites and mediate lesion resolution.

**Figure 1:**
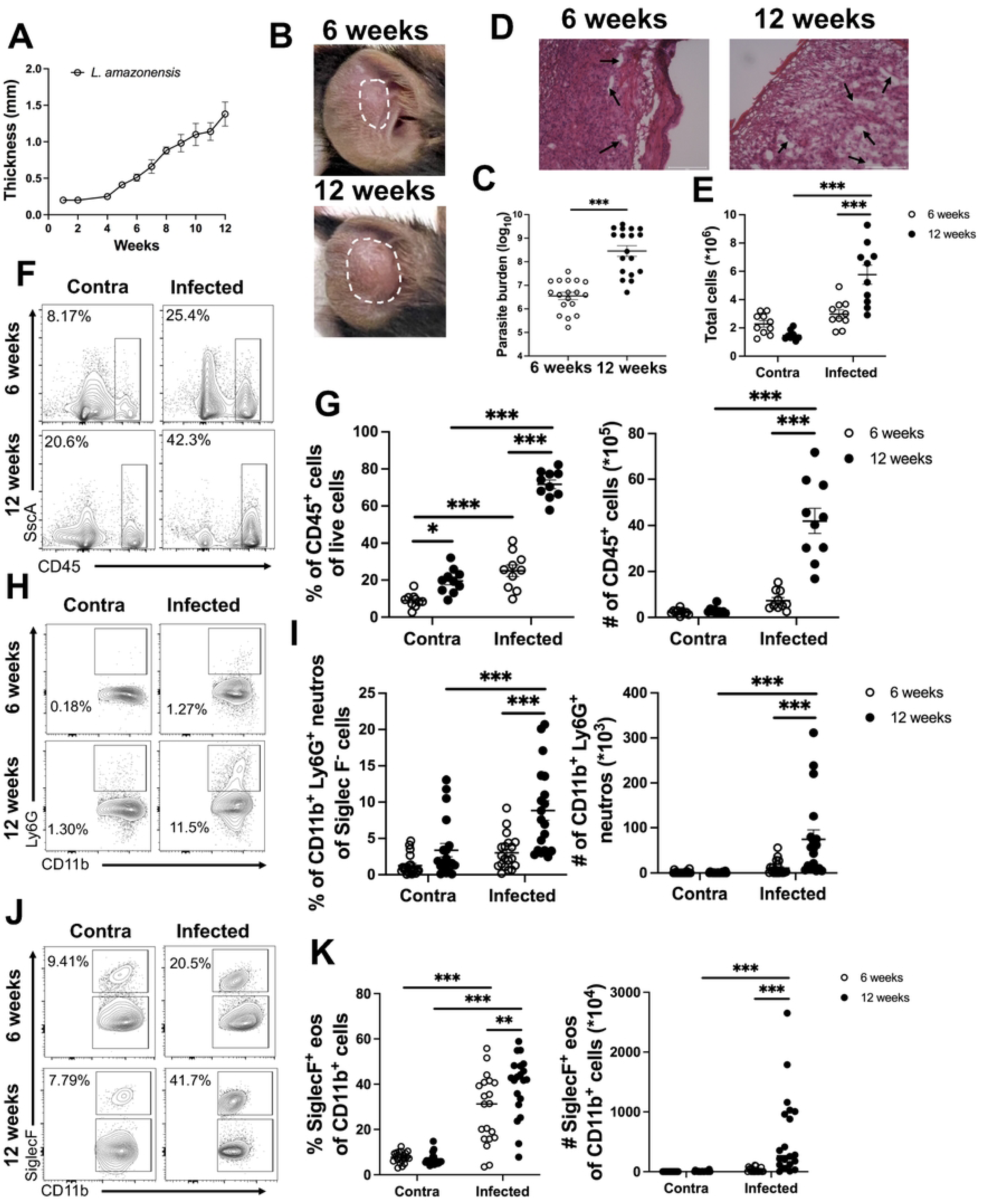
*L. amazonensis* infection leads to progressive disease and neutrophil accumulation. C57BL/6 mice were infected with 100,000 *L. amazonensis* parasites and lesions were monitored weekly. At 6 and 12 weeks post-infection (wpi), ear tissue was prepared for flow cytometry. (A) Lesion curve of ear thickness following infection with *L. amazonensis.* (B) Representative images of ear lesions in mice infected with *L. amazonensis* at 6 and 12 wpi. (C) Tissue parasite burden was determined by limiting dilution assay at 6 and 12 wpi. Data are pooled from three independent experiments, n=5 or 10 per experiment. (D) Representative images of ear sections stained with H&E from mice infected with *L. amazonensis* at 6 and 12 wpi. Immune cell infiltrate is depicted by the purple stain and white space within the tissue depicts edema. (E) Quantification of total ear cells. (F) Representative flow plots for CD45^+^ cells are shown for both a contralateral ear and infected ear at 6 and 12 wpi. (G) Quantification of the flow plots in (F) for percentage and number of CD45^+^ cells. Pre-gated on live, single cells. Data are pooled from two independent experiments where n=10. (H) Flow plots show the neutrophil population in the contralateral and infected ears at 6 and 12 wpi. (I) Quantification of (H) showing percentage and number of Ly6G^+^ neutrophils. Gated on live, single, CD45^+^, CD11b^+^, SiglecF^-^ cells. (J) Flow plots show the eosinophil population in the contralateral and infected ears at 6 and 12 wpi. (K) Quantification of (J) showing percentage and number of SiglecF^+^ eosinophils. Gated on live, single, CD45^+^, CD11b^+^, SiglecF^+^ cells. Data are pooled from three experiments where n=20. Data are shown as mean ± SEM. Significance was determined using either a student’s t-test or a two-way ANOVA paired with a Tukey’s multiple comparison test where *p<0.05 and ***p<0.001. Scale bars 100 μm.

Development of progressive lesions following *L. amazonensis* infection suggests overtime cellular infiltrate and fluid accumulate at the lesional site. To test the hypothesis that non-healing CL develops due to an accumulation of inflammatory cells and fluid, the immune cell infiltration and tissue edema in mice infected with *L. amazonensis* was evaluated at 6 and 12 weeks after parasite inoculation. H&E staining on ear sections from mice infected for 6 weeks displayed a large immune cell infiltrate and few areas of edema (Figure 1D). At 12 wpi, ear sections still contain a significant immune cell infiltrate with increased areas of edema suggesting fluid builds up in the ear from 6 to 12 wpi (Figure 1D). To further characterize the immune cell infiltrate in the ear at the site of infection, we performed flow cytometry at 6 and 12 wpi. Contralateral ear tissue from both 6 and 12 wpi was used as an uninflamed control. We found total cells in the ear increase overtime during *L. amazonensis* infection (Figure 1E). More specifically, total CD45^+^ hematopoietic cells accumulate in the lesion at 12 wpi compared to 6 wpi (Figure 1F-G). At 6 and 12 wpi similar percentages of macrophages and monocytes was detected (Figure S1A-B). In contrast, we found a significant increase in the percentage and the number of neutrophils and eosinophils at 12 wpi compared to 6 wpi (Figure 1H-K). These data suggest an accumulation of neutrophils and eosinophils at the infection site contributes to the enhanced inflammation at 12 wpi compared to 6 wpi and may promote lesion progression.

The lymphatic vasculature is tasked with taking up debris, interstitial fluid, and cells during tissue edema to return to homeostasis [32,33]. During an insult, inflammatory lymphangiogenesis allows for lymphatic remodeling, a process that expands the initial lymphatics, increasing the avenues of exit from the inflammatory site to the circulation [34]. The increased accumulation of both fluid and immune cells at the lesion site led us to hypothesize that non-healing CL is characterized by lymphatic dysregulation. To test this hypothesis, we aimed to determine if lymphatic remodeling is compromised during non-healing CL. Therefore, immunofluorescence microscopy (IF) was performed to visualize LYVE-1, a marker of LECs, and quantify LVs at both 6 and 12 wpi of *L. amazonensis* infection. The percentage of LYVE-1, relative to the region of interest (ROI) area, is used as a surrogate for lymphangiogenesis, the process of new lymphatic growth from pre-existing vessels [35]. LVs were detected at both 6 and 12 wpi (Figure 2A). Of note, LYVE-1^+^ cells were almost exclusively associated with vessels and not as individual cells in the skin, suggesting these cells are LECs and not resident LYVE-1^+^ macrophages in the skin. The percentage of LYVE-1^+^ LECs was significantly lower at 12 wpi compared to 6 wpi, even though lesions had increased in size (Figure 2B). To control for lesion size, a ROI was examined for LYVE-1^+^ LVs between time points after *L. amazonensis* infection. Ultimately, the decrease in LYVE-1^+^ LECs was associated with an overall decrease in total LVs per mm^2^ at 12 wpi compared to 6 wpi (Figure 2C). Together these data suggest non-healing CL is associated with a defect in lymphangiogenesis which is concomitant with an accumulation of immune cells and fluid (Figure 1 and 2).

**Figure 2:**
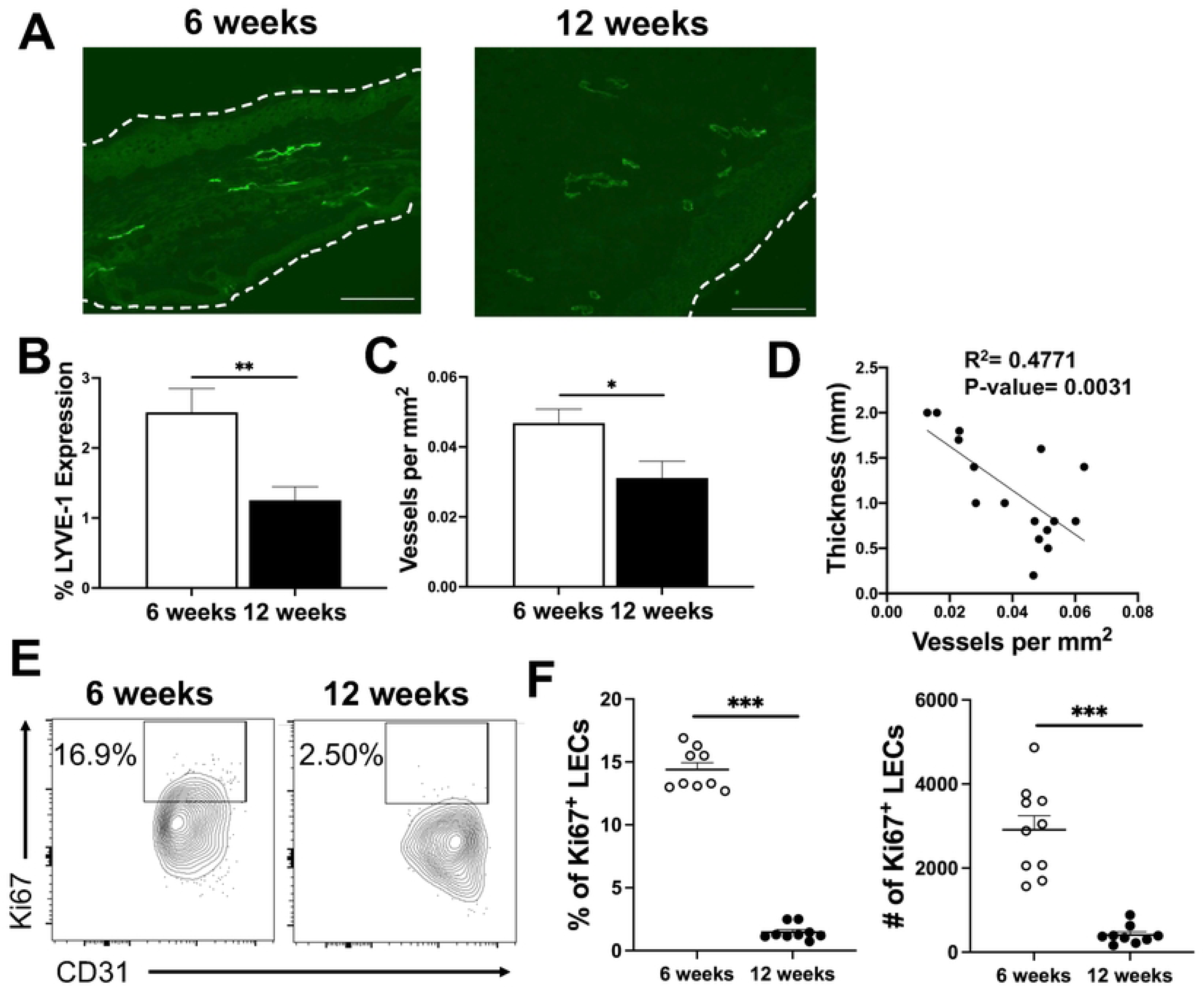
Lymphatic remodeling is compromised during *L. amazonensis* infection. Ear sections from infected mice were stained with an antibody against LYVE-1 to visualize lymphatic vessels (LVs) at 6 and 12 wpi. (A) Representative image of LVs in the ear at 6 and 12 wpi during *L. amazonensis* infection. LVs are characterized as distinct bright green structures that aggregate in linear or circular arrangements. This designation distinguishes LVs from LYVE-1^+^ macrophages. (B-C) Quantification of (A) showing the (B) percentage of LYVE-1 and (C) overall LVs per mm^2^ relative to the area of the region of interest (ROI) at 6 and 12 wpi by IF microscopy. Data are pooled from two independent experiments where n=10. (D) Correlation analysis between thickness and LVs per mm^2^. (E) Ki67 stain was used to assess LEC proliferation. (E) Representative flow plots of Ki67^+^ LECs are shown. Gated on live, single, CD31^+^, Podo^+^ cells. (F) Quantification of (E). Data are representative of three experiments. Data are shown as mean ± SEM. Significance was determined using either a student’s t-test or a two-way ANOVA paired with a Tukey’s multiple comparison test where *p<0.05 and **p<0.01. Scale bars 100 μm.

To determine if lesion severity is associated with the lymphatic vasculature, a correlation analysis of ear thickness and LV numbers was performed. Interestingly, there was an inverse correlation between lesion thickness and LV density suggesting larger lesions possess fewer LVs (Figure 2D). We hypothesized decreased lymphangiogenesis during *L. amazonensis* infection may be due to decreased LEC proliferation. To investigate this hypothesis, mice were infected with *L. amazonensis* and LEC proliferation was assessed at 6 and 12 wpi by Ki67 expression. At 6 wpi the percentage of Ki67^+^ LECs was significantly higher in infected ears compared to the contralateral ears demonstrating infection is associated with lymphatic remodeling (Figure 2E-F). However, consistent with the reduced LYVE-1 tissue positivity at 12 wpi compared to 6 wpi (Figure 2B), there was a significant decrease in the percentage and number of LECs expressing Ki67 at 12 wpi compared to 6 wpi, indicating LEC proliferation is not sustained during *L. amazonensis* infection (Figure 2E-F). These data further support the hypothesis that there are not sufficient exit routes for the fluid and recruited immune cells to leave the infection site for lesion resolution. Altogether, these findings highlight lymphangiogenesis as a new mechanism limiting disease severity in CL.

To further highlight lymphatic dysregulation during non-healing CL, we examined lymphatic remodeling using a murine model of healing CL. Intradermal infection with *L. major* parasites in C57BL/6 mice causes lesions that peak in severity around 5-6 weeks, and lesions that begin to resolve after week 8 wpi (Figure 3A). This *L. major* healing CL model contrasts *L. amazonensis* infection where lesions do not heal (Figure 1A). To investigate lymphatic remodeling during healing CL, mice were infected with 100,000 *L. major* parasites and ear tissue was taken at 6 and 12 wpi to survey lymphatic remodeling. LVs were identified at both 6 and 12 wpi, and expression of LYVE-1^+^ LECs was consistent from 6 to 12 wpi in healing CL due to *L. major* infection (Figure 3B-C). While no differences in LYVE-1 positivity were detected over time, *L. major*-infected ears contained more LVs at 12 wpi compared to 6 wpi (Figure 3D). Taken together, these data suggest expansion of the lymphatic network aids in the drainage of cells and fluid, supporting the resolution of inflammation during healing CL.

**Figure 3:**
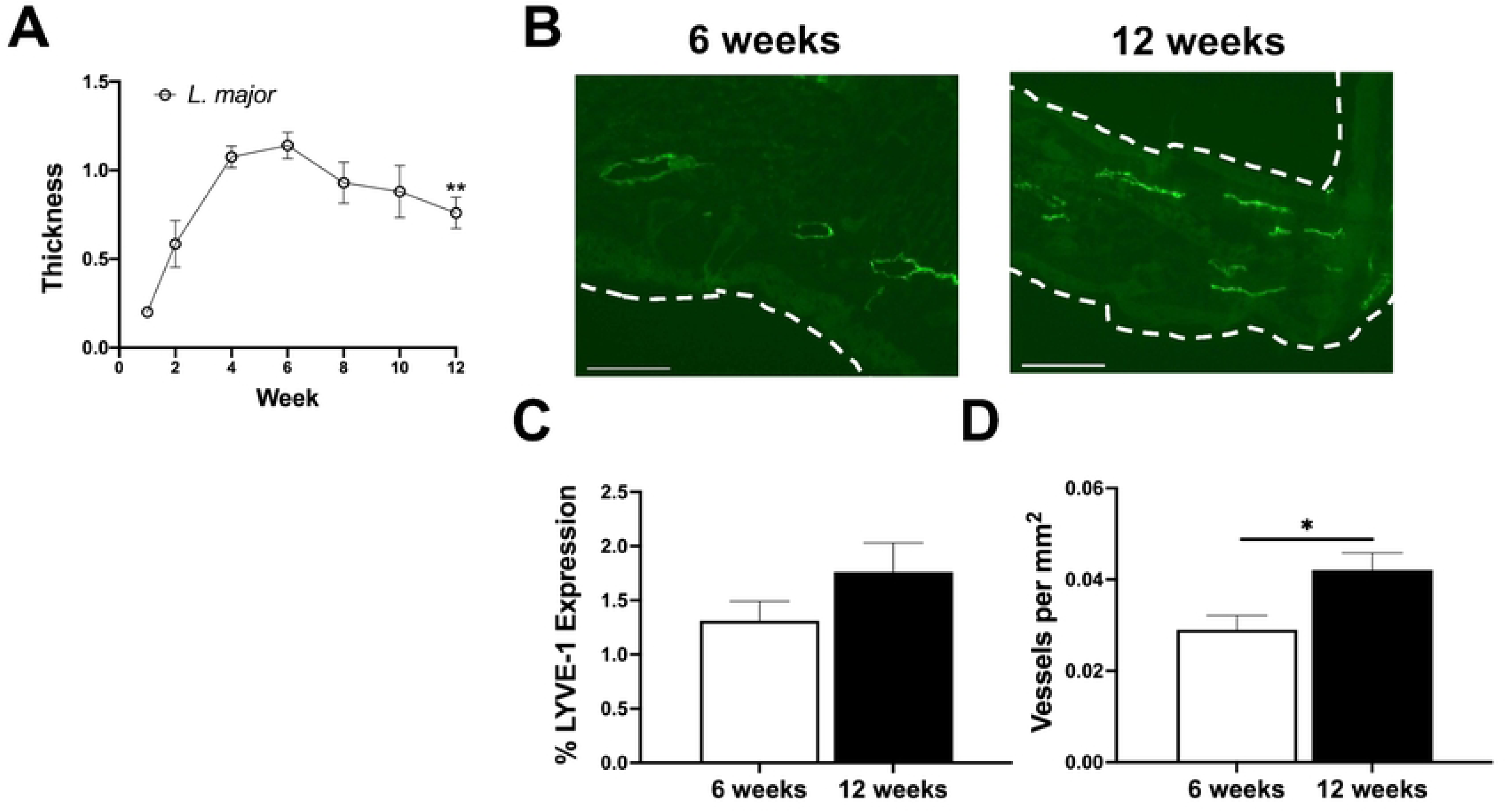
Healing CL caused by *L. major* infection possesses adequate lymphatic remodeling. C57BL/6 mice were intradermally infected with 100,000 *L. major* parasites and lesions were monitored weekly. (A) Lesion curve of ear thickness in mice infected with *L. major*. (B) At both 6 and 12 wpi ear sections were stained with anti-LYVE-1 antibody to identify LVs. Representative sections from *L. major* infected ears at 6 and 12 wpi. (C-D) Quantification of the percentage of (C) LYVE-1 and (D) LVs per mm^2^ in (B). Data are shown as mean ± SEM and significance was determined using a t-test where *p<0.05. Data is pooled from two independent experiments where n=10. Scale bars 100 μm.

Given the extensive fluid and cellular accumulation during *L. amazonensis* infection, we next aimed to explore lymphatic function. Collecting LVs located deep within the tissue actively contract to drain interstitial fluid from tissues [32,36,37]. To investigate whether lymphatic function is impaired during non-healing CL, we used a footpad infection model, where 10^4^ *L. amazonensis* parasites are injected into the footpad of C57BL/6 mice, with the contralateral footpad serving as an uninflamed control. This footpad model allows us to examine individual LVs draining the footpad to the popliteal dLN—a drainage pathway we cannot easily observe with ear dermis infections, which drain to the cervical lymph nodes. We focused on lymphatic flow and contraction rates in mice infected with *L. amazonensis* for either 3 wpi or 12 wpi. These timepoints were selected because they align with the ear dermis infection model: at 3 wpi, lesions begin to form in the footpad, while at 12 wpi, both infection models reach a chronic inflammatory state (Figure 4A). Additionally, the parasite burden is progressive in this model, increasing significantly from 3 to 12 wpi in both the local infection site as well as the dLNs (Figure 4B). At both 3 and 12 wpi, mice were injected with a FITC tracer, and drainage of individual LVs in the hindlimb were analyzed using non-invasive intravital imaging (Figure 4C-D).

**Figure 4:**
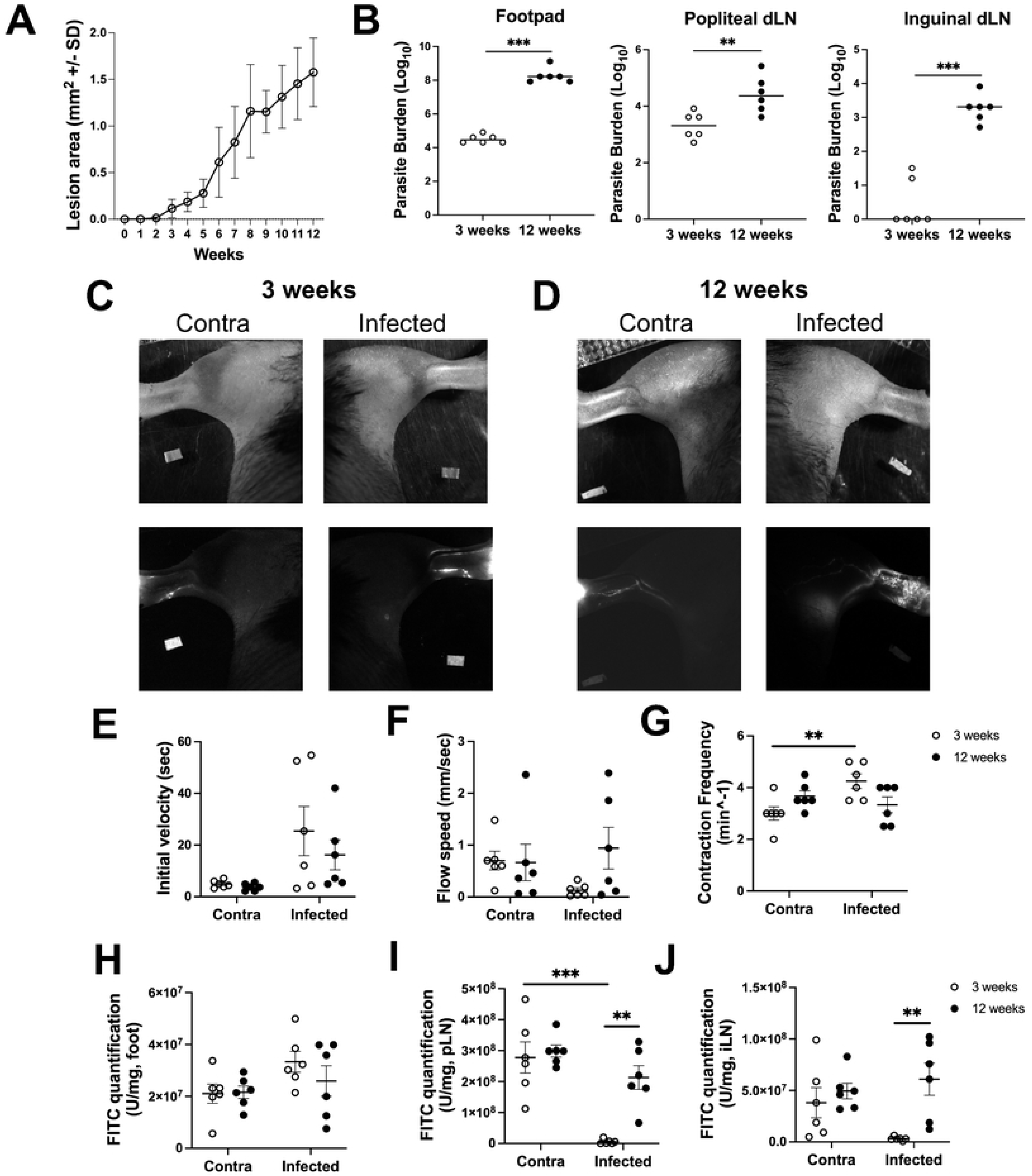
*L. amazonensis* infection compromises lymphatic function in a footpad model of infection. C57BL/6 mice were infected in the footpad of the rear hindlimb and at 3 and 12 wpi microlymphangiography was performed. (A) Lesion curve of lesion area following infection with *L. amazonensis.* (B) Tissue parasite burden was determined by limiting dilution assay at 3 and 12 wpi in the ear and the popliteal and inguinal dLN. (C-D) Representative images of the hindlimb secured for in vivo imaging and images following FITC injection tracing individual LVs at either (A) 3 wpi or (B) 12 wpi. (E) Quantification of initial velocity based on the time for the FITC signal to reach the first ROI after initial injection in (C-D). (F) Quantification of flow speed was calculated by analyzing the time for the FITC injection to go from ROI 1 to ROI 2 in (A-B). (G) Quantification of contraction frequency based on the FITC signal in (A-B). (H) Quantification of FITC signal in the foot, (I) popliteal LN, and (J) inguinal LN. Normality distribution was performed using Shapiro-Wilk Test and showed evidence of non-normality, allowing us after a visual examination of the QQ plot, to use a nonparametric test. Data are shown as mean ± SEM. Significance was determined using a two-way ANOVA paired with a Tukey’s multiple comparison test where **p<0.01 and ***p<0.001.

First, we recorded representative images of individual LVs using microlymphangiography (Figure 4C-D). These images clearly show the LVs lighting up with tracer in both the contralateral tissue and the infected footpad at 3 wpi (Figure 4C-D). Interestingly, at 12 wpi, increased vessel density is observed in the hindlimb compared to the infected hindlimb at 3 wpi. Additionally, the FITC signal is diffuse in the tissue at 12 wpi, suggesting the new LVs are somewhat leaky. Next, using non-invasive intravital imaging, lymph flow speed was assessed by defining two ROIs in the hindlimb. The first ROI was placed between the footpad and the middle of the hindlimb while the second ROI was placed further up the hindlimb closer to the dLN. By measuring the time it takes for the FITC tracer to travel from the footpad (after the initial injection) to the first ROI and followed by the time it took for the tracer to move from ROI 1 to ROI 2 we can measure lymphatic flow. To first measure lymphatic velocity, we quantified the time it took for the tracer to reach the first ROI from the initial injection site into the footpad (Figure 4E). While we did not observe significant differences in initial velocity between 3 and 12 wpi, infection at either timepoint showed a trend toward higher initial velocity compared to the contralateral control (Figure 4E). Next, we measured the time it took for the tracer to move from ROI 1 to ROI 2, which allowed for quantification of flow speed (Figure 4F). Interestingly, at 3 wpi, flow speed trended lower in the infected footpad compared to the contralateral control, suggesting that *L. amazonensis* infection impairs lymphatic function (Figure 4F). However, at 12 wpi, flow speed in the infected footpad was similar to the contralateral paw, indicating that lymphatic function was partially restored (Figure 4F). Additionally, we found that contraction frequency of the LVs was elevated at 3 wpi in the infected paw compared to the contralateral paw, indicating that, although fluid was flowing more slowly, the LVs were contracting more frequently (Figure 4G). By 12 wpi, however, no differences in contraction frequency were observed between the infected and contralateral LVs (Figure 4G). Together, these data suggest that *L. amazonensis* infection disrupts lymphatic function in the footpad at 3 wpi, but this impairment is largely resolved by 12 wpi.

To further investigate lymphatic function during footpad infection with *L. amazonensis*, we measured the amount of fluorescent tracer in the footpad, popliteal LN, and inguinal LN. By quantifying the amount of FITC in the footpad and dLNs, we can gain deeper insight into LV drainage. The tracer accumulation in the footpad was similar across infection timepoints; however, compared to the contralateral tissue, there was a trend of higher tracer accumulation in the infected footpad at 3 wpi (Figure 4H). Additionally, at 3 wpi, we observed a significant decrease in tracer accumulation in both the popliteal dLN and the inguinal dLN compared to the contralateral tissue, suggesting that reduced flow speed impairs fluid drainage to the dLNs during *L. amazonensis* infection (Figure 4I-J). At 12 wpi, however, tracer accumulation in both the popliteal and inguinal dLNs was significantly higher than at 3 wpi (Figure 4I-J). Supporting this, at 12 wpi the microlymphangiography images revealed increased LVs in the hindlimb draining the footpad to the dLNs (Figure 4D). Although more drainage routes were identified at 12 wpi, we also observed backflow of the FITC tracer within the tissue, suggesting the lymphatic drainage at 12 wpi is not fully functional. Overall, these findings suggest that *L. amazonensis* infection disrupts normal lymph flow, with potential blockage leading to increased drainage by collecting vessels to the inguinal dLN.

In addition to the major function of draining interstitial fluid, there is a growing body of evidence suggesting the lymphatic system is more than a passive conduit, playing an active role in directing leukocyte migration by expressing specific chemokines while also interacting directly with immune cells in the tissue [36,37]. Furthermore, during CL, LECs possess antigen processing machinery and MHC-II [38]. Additionally, LECs can take up and process antigen during *Leishmania* infection [38]. To further investigate lymphatic dysfunction, independent from LEC proliferation, flow cytometry was employed to analyze surface molecule expression on LECs during *L. amazonensis* infection during the initial and chronic phases of inflammation. PDL-1 and MHC-II expression were chosen for analysis because the expression of these receptors’ changes on LECs in the tumor microenvironment and these molecules also regulate tissue immune responses [39–41]. PDL-1 and MHC-II are also surrogates for LEC activation during *L. major* infection [38,42,43]. *L. amazonensis* infection significantly increased the percentage and number of LECs expressing MHC-II and PDL-1 at 6 wpi compared to LECs in the contralateral ear (Figure 5A-D). Consistent with decreased Ki67^+^ LECs at 12 wpi compared to 6 wpi, the percentage of MHC-II^+^ and PDL-1^+^ LECs significantly declined during the chronic phase of infection at 12 wpi compared to 6 wpi (Figures 5A-D). Together, these data suggest not only is lymphatic expansion dysfunctional during chronic non-healing CL (Figure 2), but other aspects of LEC function associated with regulating the immune response are dysregulated during chronic dermal inflammation.

**Figure 5:**
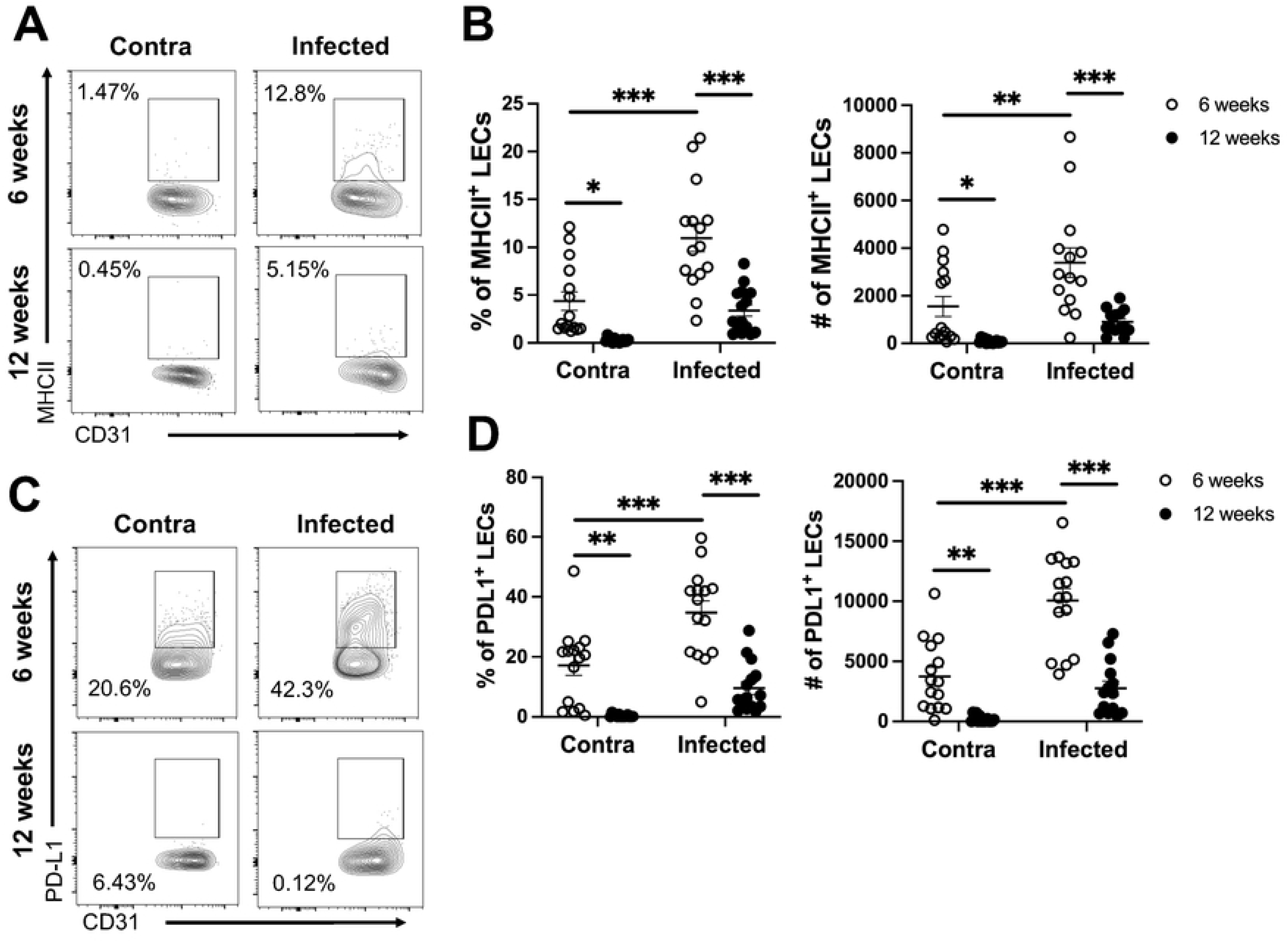
LECs do not maintain elevated MHC-II and PDL-1 expression during *L. amazonensis* infection. Mice were infected with 100,000 *L. amazonensis* parasites and surface expression on LECs was analyzed by flow cytometry. (A) MHC-II expression on LECs in the contralateral and infected ear at 6 and 12 wpi. (B) Quantification of the percentage and number of MHC-II^+^ LECs. (C) Flow plots of PDL-1^+^ expression on LECs in the contralateral and infected ear at 6 and 12 wpi. (D) Quantification of PDL-1 on LECs in the ear at 6 and 12 wpi. LECs are pre-gated on live, single, CD31^+^, Podo^+^ cells. Data are pooled from two experiments with 15 mice per group. Data are shown as mean ± SEM. Significance was determined using a two-way ANOVA paired with a Tukey’s multiple comparison test where *p<0.05 **p<0.01, and ***p<0.001.

The inflammatory microenvironment directly dictates LEC function. Specifically, type 2 cytokines inhibit lymphangiogenesis [44]. We sought to determine the environmental cues present during non-healing CL that contribute to lymphatic dysregulation [36,44–47]. We analyzed levels of cytokines at the site of infection including IFNγ, IL-10, IL-17, IL-4, and IL-13 by quantitative real-time PCR. The expression of IFNγ, IL-10, IL-17, and IL-13 were similar between 6 wpi and 12 wpi, however IL-4 expression was significantly increased at 12 wpi compared to 6 wpi suggesting a shift in the pro-inflammatory milieu towards a mixed Th1/Th2 immune response during the chronic phase of *L. amazonensis* infection (Figure 6A). Due to elevated IL-4 at 12 wpi, macrophage activation status, as a surrogate for phenotypic changes in immune cells in the lesions, was analyzed. The percentage and number of macrophages and inflammatory monocytes (iMonos) expressing Arg-1, a marker of Th2 immune responses, significantly increased from 6 to 12 wpi (Figure 6B-D). Although M2 macrophages are associated with lymphangiogenesis to promote wound healing, only low levels of vascular endothelial growth factor-C (VEGF-C) were found at both 6 and 12 wpi (Figure 6E), suggesting Arg-1^+^ macrophages are not inducing healing, consistent with lesion sizes at 12 wpi [48]. Of note, an increase in the percentage and number of activated CD44^+^CD4^+^ T cells producing IFNγ and TNF-α was detected at 12 wpi compared to 6 wpi in the dLN (Figure S2A-D). However, the elevated CD44^+^CD4^+^ T cells in the dLN producing IFNγ and TNF-α does not correlate with IFNγ expression in the ear, or differences in parasite burdens (Figure S2). Together this data suggests the chronic phase of *L. amazonensis* infection is associated with a mixed Th1/Th2 environment that may contribute to LEC dysregulation.

**Figure 6:**
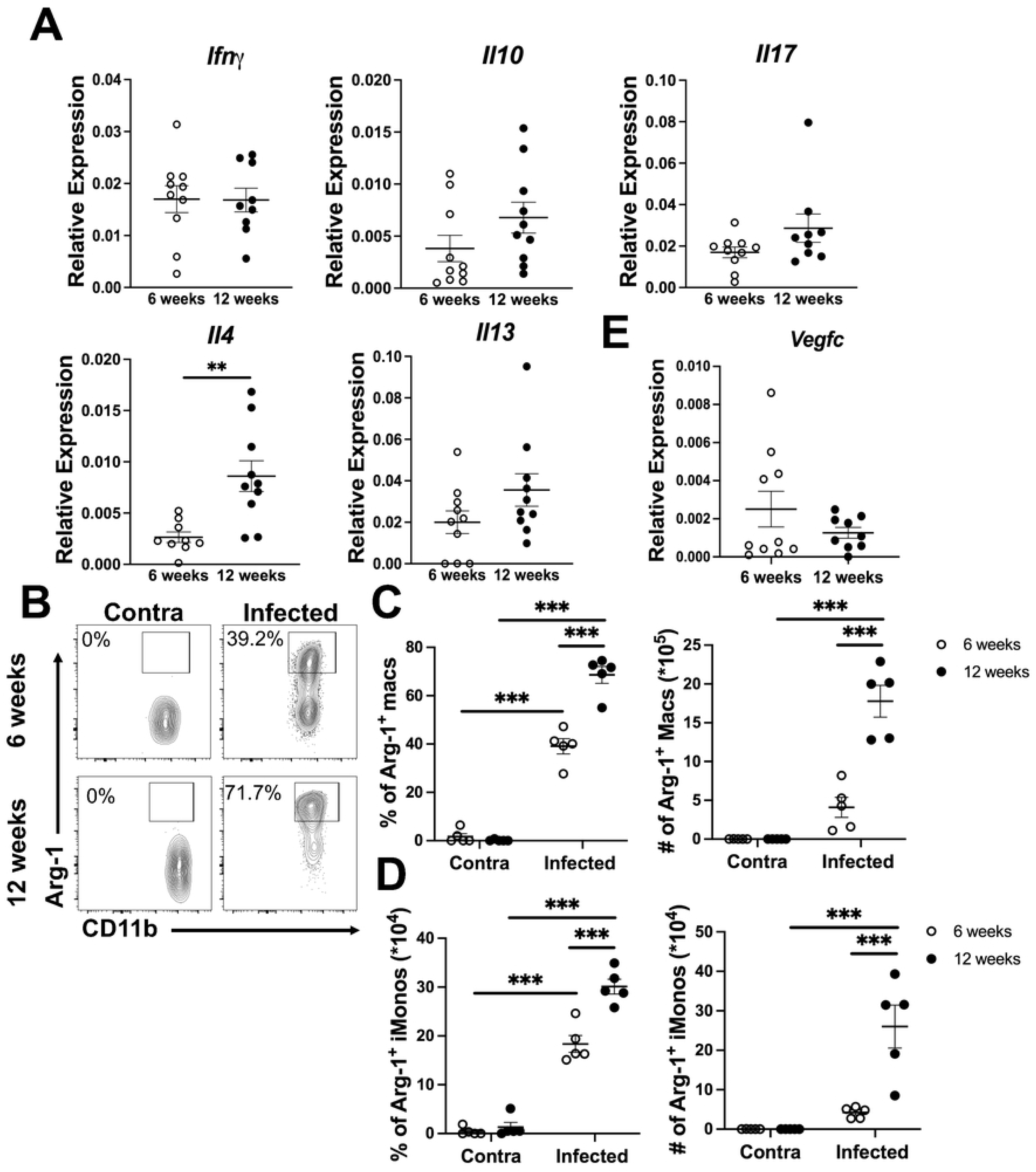
The chronic phase of *L. amazonensis* infection shifts towards a Th2 dominant microenvironment. Mice were infected with 100,000 *L. amazonensis* parasites and real-time PCR was performed on ear tissue at 6 and 12 wpi. (A) Relative expression of cytokines in the ear at 6 wpi and at 12 wpi. (B) Flow cytometric analysis was performed on ear tissue from 6 and 12 wpi. Representative flow plots of Arg-1^+^ macrophages are shown. Gated on live, single, CD45^+^, CD11b^+^, CD64^+^, Ly6G^-^ cells. (C) Quantification of (B) shows percentage and number of ARG-1^+^ macrophages at 6 and 12 wpi in the infected and contralateral ear. (D) Quantification of percentage and numbers of ARG-1^+^ monocytes at 6 and 12 wpi from ears. Gated on live, single, CD45^+^, CD11b^+^, Ly6C^+^, Ly6G^-^ cells. (E) Relative expression of VEGF-C as determined via real-time PCR. Real-time PCR data are pooled from two experiments with 10 mice per group. Data are shown as mean ± SEM. Significance was determined using either a student’s t-test or a two-way ANOVA paired with a Tukey’s multiple comparison test where **p<0.01 and ***p<0.001.

To explore the therapeutic implications of targeting dysregulated lymphatic remodeling during chronic inflammation caused by *L. amazonensis* infection, VEGF-C was administered by adenoviral delivery as previously described [49,50]. Adenoviral VEGF-C expands existing lymphatics through targeting of VEGFR-3 on LECs [50–53]. Mice were infected with *L. amazonensis* and at 2 and 7 wpi, adenoviral VEGF-C or control virus was administered. To validate the effects of adenoviral VEGF-C on the dermal lymphatics, ear sections from mice infected with *L. amazonensis* and treated with either control or VEGF-C adenovirus were taken and stained with an antibody against LYVE-1 (Figure 7A). Adenoviral delivery of VEGF-C induced lymphatic remodeling, specifically lymphangiectasia, where mice treated with VEGF-C show higher levels of LV dilation compared to mice treated with control adenovirus (Figure 7B). However, there were no differences in percentage of LYVE-1 or total vessels per mm^2^ between mice treated with control or VEGF-C adenovirus (Figure 7C-D). Along with an increase in LV dilation, mice treated with VEGF-C possessed a higher percentage of Ki67^+^ proliferating LECs compared to mice treated with control virus (Figure 7E). Ultimately, lymphatic remodeling induced by VEGF-C administration led to significantly decreased lesional volume in mice compared to those treated with an empty vector control virus at 9 wpi (Figure 7F). Importantly, adenoviral VEGF-C did not alter the parasite burden compared to controls (Figure 7G). Taken together, these data show VEGF-C targets the lymphatics leading to a reduction in lesion sizes without altering parasite burdens.

**Figure 7:**
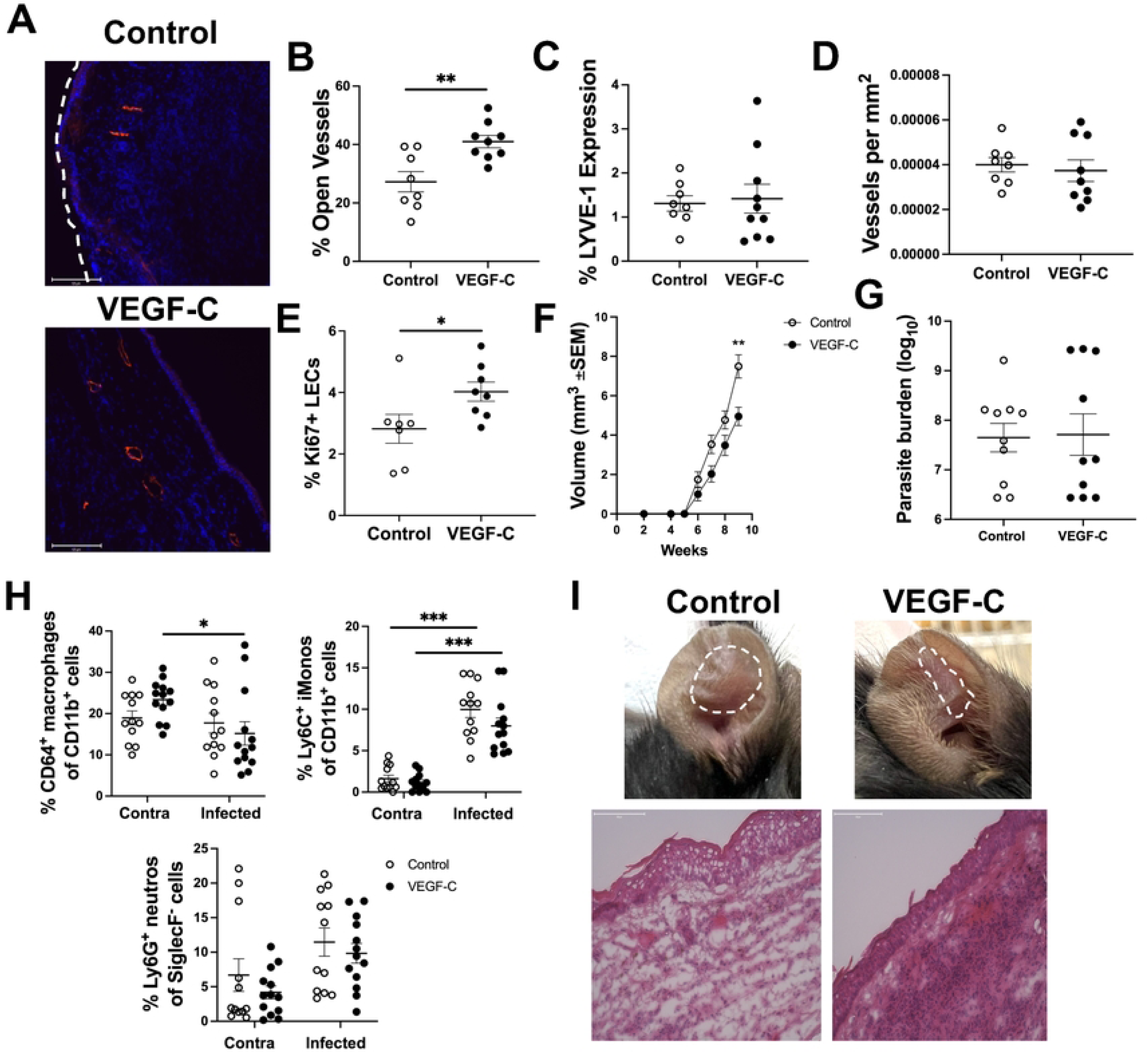
Adenoviral delivery of VEGF-C decreases disease severity during *L. amazonensis* infection. At 2 and 7 wpi with *L. amazonensis*, control or VEGF-C adenovirus was administered, and disease severity was measured weekly. (A) Ears from infected mice given control or VEGF-C adenovirus were stained with an antibody against LYVE-1 at 9 wpi. (B) Quantification of (A) showing percentage of dilated LVs, (C) percentage of LYVE-1, and (D) total vessels per mm^2^. (E) Ki67 stain was utilized to assess LEC proliferation in control- or VEGF-C-treated mice at 9 wpi. Gated on live, single, CD31^+^, Podo^+^ cells. (F) Lesion volume from mice given control or VEGF-C adenovirus. (G) Parasite burden in the ear was determined using a limiting dilution assay at 9 wpi (n=10 per group). Data are pooled between 2-3 experiments and shown as mean ± SEM. (H) Percentages of CD64^+^ macrophages (gated on live, single, CD45^+^, CD11b^+^, Ly6G^-^), Ly6C^+^ monocytes (gated on live, single, CD45^+^, CD11b^+^, Ly6G^-^), and Ly6G^+^ neutrophils (gated on live, single, CD45^+^, CD11b^+^, SiglecF^-^) during infection with *L. amazonensis* and following treatment with control or VEGF-C adenovirus. (I) Representative images of lesions or representative H&E staining of ear sections from mice infected with *L. amazonensis* and treated with either control or VEGF-C adenovirus. Immune cell infiltrate is depicted by the purple stain and white space within the tissue depicts edema. The flow cytometric Ki67 analysis is representative of 1 experiment. Significance was determined using either a student’s t-test or a two-way ANOVA paired with a Tukey’s multiple comparison test where *p<0.05 and **p<0.01. Scale bars 125 μm.

We hypothesize VEGF-C treatment enhances fluid and cellular drainage from the ear. Interestingly, VEGF-C treatment did not alter the percentages of key cell types involved in the anti-parasitic response, such as macrophages, iMonos, and neutrophils, when compared to control adenovirus-treated mice (Figure 7H). This suggests that VEGF-C’s primary effect is on fluid drainage rather than modulation of the anti-parasitic immune response, which remains intact during *L. amazonensis* infection (Figure S3). In support of VEGF-C alleviating fluid buildup, H&E stain on sections from mice treated with control or VEGF-C adenovirus demonstrate that while VEGF-C sections present with less edema in comparison to control sections, the cellular infiltrate is still present (Figure 7I).

To confirm, that VEGF-C decreases lesion severity without effecting anti-parasitic programs, we investigated dLN CD4^+^ and CD8^+^ T cells in mice infected with *L. amazonensis* and treated with VEGF-C or adenovirus controls, given T cell responses are key to parasite control in CL. Despite differences in lesion severity, there were no detectable differences in the dLN populations in VEGF-C-treated mice versus control-treated mice. Specifically, no differences were detected in the CD4^+^ or CD8^+^ compartments including CD4^+^ and CD8^+^ cells as well as activated CD44^+^CD4^+^ or CD44^+^CD8^+^ cells (Figure S3A-D). Additionally, intracellular cytokine analysis indicated similar IFNγ and TNF-α production from activated CD4^+^ and CD8^+^ cells (Figure S3E-F and data not shown). Additionally,while *L. amazonensis* infection typically induces a mixed Th1/Th2 microenvironment in the chronic phase (Figure 6), we did not observe any changes in the Th1-associated cytokine IFNγ or other cytokines implicated in the response to *L. amazonensis* parasites such as IL-17, or IL-10 in response to VEGF-C treatment (Figure S3G). Together, these data show VEGF-C impacts lymphatic remodeling during skin infection by increasing dermal LEC proliferation and LV dilation, supporting fluid drainage to the dLN compared to control-treated mice. Importantly, these data demonstrate targeting lymphatic remodeling by VEGF-C reduces disease severity without impairing anti-parasitic immune responses.

## Discussion

Understanding factors limiting healing during CL is crucial in developing host-targeted therapeutics for patients resistant to conventional anti-leishmanial treatments. Here, for the first time, we show lymphatic insufficiency as a mechanism for exacerbated immunopathology during non-healing CL. During the progression from acute to chronic *L. amazonensis* infection, we report fluid and neutrophils accumulate in *L. amazonensis* lesions which is associated with less new lymphatic growth and defects in lymphatic function in response to inflammation (Figure 1, 2 and 4). Additionally, although IFN-γ levels are consistent at 6 and 12 wpi, the LEC activation signature is dysregulated during chronic inflammation (Figure 5). This contrasts with what we see during *L. major* infection and healing CL where lymphatic remodeling is sustained until inflammation resolves (Figure 3). We hypothesize reduced lymphangiogenesis and decreased total lymphatics hinders cellular and fluid escape from the lesion perpetuating inflammation and eliciting chronic disease. Ultimately, this study demonstrates the lymphatics as a therapeutic target during non-healing CL to alleviate disease following *L. amazonensis* infection.

This study demonstrates that infection-induced changes in the lymphatic system significantly impact chronic inflammation. While lymphatic dysfunction clearly arises following *L. amazonensis* infection, the underlying mechanisms remain unclear. A key distinction between immune responses to *L. amazonensis* and *L. major* infections is the dominance of IFN-γ during *L. major* infection, which subsides after parasite control. In contrast, *L. amazonensis* infection is characterized by a mixed Th1 and Th2 response, with sustained levels of both IL-4 and IFN-γ even at 12 wpi. LECs have receptors to respond to these cytokines, suggesting that the persistent expression of IFN-γ and increased IL-4 during *L. amazonensis* infection may suppress new lymphatic growth [44,54].

In vitro and in vivo evidence indicates that both IFN-γ and IL-4 inhibit LEC proliferation and tube formation [44,54–58]. For example, Katura *et al.* found that areas rich in IFN-γ-producing CD4+ T cells in tumor-dLNs failed to expand LVs; however, deleting IFN-γ from lymphocytes significantly increased LV density [57]. Additionally, targeting the IFN-γ receptor on LECs in a metastatic tumor model led to increased LYVE-1 expression and reduced tumor burdens [59]. Zampbell *et al.* proposed that impaired lymphatic function results from the dysregulation of pro- and anti-lymphangiogenic mediators [58]. Neutralizing IFN-γ enhanced inflammatory lymphangiogenesis regardless of VEGF-A and VEGF-C levels, while blocking Th2 cytokines, IL-4 and IL-13, reduced lymphedema and improved lymphatic function [58]. Finally, IL-4 and IL-13 blockade in the cornea specifically resulted in heightened lymphangiogenesis, but not angiogenesis [44]. Collectively, these findings suggest that Th1 and Th2 cytokines negatively regulate lymphangiogenesis and may contribute to lymphatic dysfunction during chronic *L. amazonensis* infection. While challenging because Th1 and Th2 cytokines regulate parasite control, future research will investigate the impact of Th1 and Th2 cytokines on the lymphatic endothelium during *L. amazonensis* infection.

This report is the first to show that lymphatic function is disrupted during *L. amazonensis* infection, in addition to defects in lymphangiogenesis. Following infection in the footpad, flow speed decreases and contraction frequency increases suggesting compensation by the LVs. At 3 wpi, minimal FITC tracer drainage reaches the popliteal or inguinal dLNs, indicating compromised lymphatic function. However, by 12 wpi, these defects are largely resolved: flow speed normalizes and tracer reaches both the popliteal and inguinal dLNs. Unexpectedly, we also observe an expansion of the lymphatic network in the hindlimb by 12 wpi. Intravital imaging shows elevated LVs draining from the foot to the dLNs. We believe this represents the formation of new collecting vessels, which may redirect drainage to the inguinal LN, compensating for the impaired drainage to the popliteal dLN observed at 3 wpi. However, at 12 wpi some of the FITC tracer was not associated with individual LVs suggesting there is backflow occurring during chronic *L. amazonensis* infection. Interestingly, the formation of “collateral” LVs has been observed following ligation of collecting LVs in the hindlimb which redirect fluid drainage [60].

Our findings show exogenous VEGF-C improves lesion sizes during CL. Targeting of VEGF-C is an active area of research in treatment for secondary lymphedema and is currently undergoing phase 1 clinical trials [61–64]. Specifically, lymphatic gene therapy, involving the administration of VEGF-C or VEGF-D, has promising disease outcomes by selectively targeting the lymphatics, without affecting the blood vasculature [65]. Moreover, stimulation of lymphangiogenesis through VEGF-C administration has been demonstrated in models of cutaneous inflammation [53,66,67]. Using repeated exposure to oxazolone as a model of skin inflammation, Huggenberger *et. al* found administration of VEGF-C limited dermal inflammation through expansion of the lymphatics resulting in decreased cellular infiltrate in the skin and overall, more intact epidermal architecture [51]. We predicted VEGF-C would increase total LVs in the ear compared to control treated mice. While we did detect a slight, yet seemingly temporary, increase in the Ki67^+^ LECs following VEGF-C administration, this did not result in an overall increase in LV density (Figure 6E). We anticipate we did not detect an increase in total LVs because VEGF-C administration in the context of *L. amazonensis* infection appears to transiently change the morphology of LVs.

Our findings support VEGF-C as a promising therapeutic target for lymphatic dysfunction. We provide evidence that VEGF-C enhances LEC proliferation and LV dilation, a process known as lymphangiectasia, which significantly improves outcomes for non-healing CL. VEGF-C therapy has been reported to instigate LV dilation which ultimately improved CSF out of the brain [68]. We speculate that interstitial pressure gradients facilitate LV dilation, enabling fluid to escape from lesions during VEGF-C administration. Interestingly, we have recorded increases in LV dilation following *L. amazonensis* infection compared to naïve mice, but we saw no changes in LV dilation when progressing from acute to chronic inflammation. This suggests that, in the context of *L. amazonensis* infection, LV dilation may be compromised.

Since we did not observe a decrease in cell numbers in the ears of mice treated with VEGF-C adenovirus compared to controls, we hypothesize that VEGF-C therapy alleviates fluid buildup, thus improving chronic inflammation. Importantly, VEGF-C prophylaxis promotes CSF drainage in a murine model of ischemic stroke resulting in better neurological outcomes [69]. The protection of VEGF-C during ischemic strokes is lost upon cauterization of the afferent lymphatics of the cervical dLN where CSF can no longer drain [69]. We propose that fluid and immune cells enter the initial lymphatics via different mechanisms: fluid flows through button junctions between LECs, while immune cells utilize receptor-ligand interactions with adhesion molecules on the lymphatic endothelium. As LV dilation occurs, these junctions expand, allowing increased fluid entry. Additionally, phagocytes co-cultured with *Leishmania* parasites exhibit reduced adhesion, impairing their entry into the dLN [70]. This suggests that regulated entry differences may explain the lack of changes in cellular populations following VEGF-C administration. Future research should explore fluid drainage dynamics after VEGF-C treatment and adhesion molecule expression on immune cells to clarify how VEGF-C reduces immunopathology.

Ultimately, this work adds to the expanding body of literature highlighting the lymphatic system as a crucial therapeutic target in managing infectious diseases. One of the most notable examples of infection-induced lymphatic dysregulation occurs during lymphatic filariases. Parasites target the lymphatics, resulting in dysfunctional lymph flow leading to the development of lymphedema and altered immune responses in the tissue [71,72]. Similarly, bacterial infections, such as those caused by methicillin-resistant *S. aureus,* also lead to lymphatic dysfunction [73]. In response to cutaneous vaccinia virus infection by scarification, LECs tighten their connections, restricting fluid transport and sequestering the virus to the skin, preventing metastasis [74]. Finally, during infection with *Toxoplasma. gondii*, a neurotropic parasite, drainage of cerebral spinal fluid through meningeal lymphatics is significantly reduced in infected mice compared to naïve mice [49]. Importantly, VEGF-C administration-initiated drainage of cerebral spinal fluid, but did not change *Toxoplasma*-specific T cell responses in the dLN like what is seen in the dLN during *L. amazonensis* infection [49]. Understanding the mechanisms by which infections impact lymphatic dysregulation is crucial for elucidating disease pathogenesis and will inform the design of therapeutic strategies targeting the lymphatics in infectious diseases.

## Material and Methods

### Mice

Female C57BL/6NCr mice were purchased from the National Cancer Institute. Mice were used for experiments at 6 to 8 weeks of age and were housed under pathogen-free conditions at the University of Arkansas for Medical Sciences. All procedures were approved by the UAMS or Calgary IACUC and performed in accordance with institutional guidelines.

### Parasites

*Leishmania amazonensis* (MHOM/BR/00/LTB0016) and *L. major* (WHO/MHOM/IL/80/Friedlin) parasites were utilized for the ear dermis infection model while *L. amazonensis* (IFLA/BR/67/PH8) were used for the footpad infection model. Parasites were grown in vitro in Schneider’s Drosophila medium (Gibco) supplemented with 20% heat-inactivated fetal bovine serum (FBS, Invitrogen), 2 mM L-glutamine (Sigma), 100 U/mL penicillin, and 100 µg/mL streptomycin (Sigma). Metacyclic promastigotes from 4-5-day old parasite cultures were isolated by Ficoll density gradient centrifugation (Sigma) [75].

### In vivo infections

For ear dermis infections, 100,000 parasites were intradermally injected into the right ear in a volume of 10 µL of phosphate-buffered saline (PBS) (Gibco). For footpad infections, 10,000 parasites were utilized for the infection dose. The contralateral left ear or footpad was left uninfected, serving as an uninflamed control. Ear thickness and lesion diameters were recorded weekly with electronic calipers and lesion volume was calculated. Ears were digested enzymatically for 90 minutes at 37°C in 0.25 mg/mL liberase (Roche) with 10 µg/mL DNase I (Sigma) in RPMI 1640 (Gibco). Tissue parasite burden was determined using limiting dilution assays [76].

### Flow cytometry

Ear tissue was enzymatically digested and processed for flow cytometric analysis of dermal cells from the ears. For dLNs, cells were stimulated with brefeldin A (BFA; 1:1,000; eBioscience), monensin (1:1,000; eBioscience), PMA (100 ng/mL; Abcam), and ionomycin (1 mg/mL; Sigma) for 4 h before viability and surface staining. To determine cellular viability, cells were incubated with a Zombie Aqua viability dye (Biolegend). Fc receptors were blocked with 2.4G2 anti-mouse CD16/CD32 (Invitrogen or BioXCell) and 0.2% rat IgG (BioXCell). Cells were surfaced stained with antibodies against CD45 BV650 (BD Horizon) or AF700 (eBioscience), CD11b BV750 (Biolegend) or BV605 (Biolegend), Siglec-F BB515 (BD Horizon), MHC-II APC-Fire 750 (Biolegend), CD11c SB550 (Biolegend), CD64 BV421 (Biolegend) or BV711 (Biolegend), Ly6C PercpCy5.5 (Invitrogen), Ly6G AF700 (Biolegend), CD3e SB436 (Invitrogen) or APC ef780 (eBiosciences), CD4 PE-Cy5 (Tonbo Biosciences) or BV650 (BD), CD8β BV786 (BD Optibuild) or PE-TexasRed (Invitrogen), CD31 ef450 (Invitrogen), Podoplanin AF647 (Biolegend), CD44 ef450 (eBioscience), PDL-1 BV650 (Biolegend). Intracellular stain was done using the Foxp3 Transcription Factor Staining Buffer kit (Invitrogen). Intracellular molecules and cytokines were stained with antibodies against IL-17 AF488 (eBioscience), IFNγ Pecy7 (eBioscience), Ki67 Pecy7 (Invitrogen), and Arg-1 PE (Invitrogen). Cell events were acquired on a Cytek Northern Lights (Cytek) and analyzed using FlowJo (Tree star).

### Histology and Immunofluorescence microscopy

Infected ears from mice infected with *L. amazonensis* or *L. major* were frozen in Tissue-Tek OCT (Sakura). For histology, sections were subjected to hematoxylin and eosin (H&E) stain. For Immunofluorescence microscopy, tissue sections were fixed with 3% formaldehyde, blocked with 10% goat serum (Sigma) for 1 h, stained with rabbit anti-LYVE-1 antibody (AngioBio) overnight, and followed by goat anti-rabbit AF488 secondary antibody (Life Technologies) or goat anti-rabbit AF647 secondary antibody (Invitrogen) for 1 h. Ear sections were mounted using Prolong Gold with DAPI (Invitrogen). Sections were imaged using an EVOS Fl Auto 2 microscopy (Invitrogen). Fluorescent images were analyzed using Volocity (Quorum Technologies Inc.) In brief, the region of interest (ROI) tool was used to draw an ROI around the section for normalization excluding: 1) the epidermis (due to non-specific binding of the secondary antibody), and 2) folded areas due to increased background brightness relative to flat tissue. To determine LV density, individual LYVE-1^+^ LVs were counted within the ROI and the number of LYVE-1^+^ LVs was divided by total area of the ROI. The percent of LYVE-1^+^ expression was measured by calculating the area of LYVE-1^+^ pixels and dividing the area of pixels by the total area of the ROI and multiplying by 100. The percent of dilated vessels was determined by counting all the vessels within the ROI that possessed a visible lumen (considered dilated) divided by the number of total vessels (both dilated and not) within the ROI. A lumen above an area of 4.2 was considered dilated.

### mRNA extraction and real time PCR

mRNA was extracted with the Qiagen RNeasy Mini Kit 250 (Qiagen). RNA was reverse-transcribed with the High-Capacity cDNA reverse transcription kit (Applied Biosystems). Quantitative real-time PCR was performed using SYBR green PCR Master Mix and a QuantStudio 6 Flex real-time PCR system (Life Technologies). Mouse primer sequences were selected from the PrimerBank (http://pga.mgh.harvard.edu/primerbank/): *Ifnγ* (forward 5’-GACTGTGATTGCGGGGTTTGT-3’ and reverse 5’-GGCCCGGAGTGTAGACATCT-3’), *Il4* (forward 5’-ATGGAGCTGCAGAGACTCTT-3’ and reverse 5’-AAAGCATCGGTGGCTCAGTAC-3’), *Il10* (forward 5’-TGTCCAGCTGGTCCTTTGTT-3’), *Il17a* (forward 5’-TCAGCGTGTCCAAACACTGAG-3’ and reverse 5’-CGCCAAGGGAGTTAAAGACTT-3’), *Il13* (forward 5’-TGAGCAACATCACACAAGACC-3’ and reverse 5’-GGCCTTGCGGTTACAGAGG), *Ccl11* (forward 5’-GAATCACCAACAACAGATGCAC-3’ and reverse 5’-ATCCTGGACCCACTTCTTCTT-3’), *Vegfc* (forward 5’-GAGGTCAAGGTCTTTGAAGGC-3’ and reverse 5’-CTGTCCTGGTATTGAGGGTGG-3’), and *RpsII* (forward 5’-CGTGACGAAGATGAAGATGC-3’ and reverse 5’-GCACATTGAATCGCACAGTC-3’).

### Non-invasive intravital imaging

After 3 weeks of *L. amazonensis* infection the mouse was anesthetized with an intraperitoneal injection of 100 mg/kg ketamine plus 10 mg/kg xylazine, and the veet hair removal cream was used to shave their legs. The mouse was then positioned face down on a flat surface, with the paw secured to maintain the leg in a stable orientation suitable for *in vivo* imaging. This setup allowed for the observation of velocity, lymph flow speed, and contraction frequency in the popliteal LV to the popliteal LN. After 30 min of equilibration, the non-invasive real-time imaging, using a stereomicroscope (Leica AF 6000, M165FC), was started 1 min before the intradermal injection (30-gauge needle) of 10 µl 2% fluorescein isothiocyanate (FITC)-dextran diluted in phosphate-buffered saline (PBS) into the mouse footpad, as previously described [77]. The fluorescent excitation was captured with the GFP laser of the steromicroscope camera (Leica-DFC9000GtC-VSC05241) 2 fps for 30 min (total 3600 frames), no binning, magnification 0.73x, and gain 2 were unchanged during the experiment. Two regions of interest (ROIs) of fixed size were placed over the pLV 4 mm apart to assess the first 6 min of flow speed of the lymph marked with FITC. The velocity was calculated by measuring the time elapsed from the initial injection until the tracer reached ROI 1 (seconds), and the flow speed was calculated by the ROI 2 (seconds) – ROI 1 (seconds) / distance (mm) between ROI 1 and ROI 2. In the last min of video capture, a fixed size ROI was placed to measure the contraction frequency of the vessel visualized by the FITC signal.

### Evaluation of FITC drainage to the dLNs

Drainage to the popliteal dLN (pdLN) and inguinal dLN (idLN) was assessed 24 h after FITC-dextran injection in the footpad as previously described. Briefly, either the pdLN or the idLN was collected, weighed, and homogenized. 100 μl of the homogenate was placed in duplicates in the wells of the multilabel plate reader (Perkin Elmer Victor X3, model 2030-0030, USA) and fluorescence intensity of duplicate samples of each well measured at an excitation wavelength of 485 nm and an emission wavelength of 535 nm.

### VEGF-C adenovirus

Adenovirus CMV VEGFC-GFP (lot #33803) and control VQ adenovirus empty-GFP (lot #24903) were purchased from Viraquest. An injection dose of 5.00E + 8 adenovirus was administered to mice through retro-orbital injection in a volume of 50 µL of PBS at 2 and 7 weeks after inoculation with *L. amazonensis* in the ear.

### Statistics

All data were analyzed for statistical significance using GraphPad Prism 9. Statistical significance was determined using either a two-tailed Student’s unpaired t-test or a two-way anova with a Tukey’s multiple comparison test. Outliers were identified by a Grubb’s outlier test and removed.

## Acknowledgments

This work was supported by the Center for Microbial Pathogenesis and Host Inflammatory Responses (funded by NIH NIGMS Centers of Biomedical Research Excellence Grant P20-GM103625). This publication was also supported in part by funds provided by the National Center for Advancing Translational Sciences of the NIH under awards TL1 TR003109 and UL1 TR003107 for the Systems Pharmacology and Therapeutics (SPaT) NIH T32 training grant GM106999 to Lucy Fry. The content is solely the responsibility of the authors and does not necessarily represent the official views of the NIH. The funders had no role in study design, data analysis, decision to publish or preparation of the manuscript.

## FIGURE LEGENDS

**Supplemental Figure 1: Myeloid-derived cell populations are consistent in the ear at 6 and 12 wpi during *L. amazonensis* infection.** (A) Gating strategy for flow cytometric analysis on immune and non-immune compartments during *L. amazonensis* infection. (B) Quantification of percentages of CD64^+^ macrophages (gated on live, single, CD45^+^, CD11b^+^, CD64^+^, Ly6G^-^) and Ly6C^+^ monocytes (gated on live, single, CD45^+^, CD11b^+^, Ly6C^+^, Ly6G^-^). Population gates are assigned based on fluorescence minus one (FMO) controls. Data are shown as mean ± SEM and significance was determined using a student’s t-test.

**Supplemental Figure 2: The immune response in the dLN is exacerbated late during *L. amazonensis* infection.** Mice were infected with 100,000 *L. amazonensis* parasites and dLNs were collected at 6 and 12 wpi. (A) Representative flow plots of CD45^+^CD3^+^CD4^+^CD44^+^ IFNγ^+^ populations in the contralateral or infected ear at 6 and 12 wpi. (B) Quantification of (A). (C) Representative flow plots of CD45^+^CD3^+^CD4^+^TNF-α^+^ cells in the contralateral and infected ears at 6 and 12 wpi. (D) Quantification of (C). Data are show from one experiment with 5 mice per group. Data are shown as mean ± SEM. Significance was determined using a two-way ANOVA paired with a Tukey’s multiple comparison test where ***p<0.001.

**Supplemental Figure 3: dLN immune responses are not altered following adenoviral VEGF-C administration during *L. amazonensis* infection.** Two independent experiments were performed analyzing the dLN from the infected and contralateral ear during *L. amazonensis* infection and during treatment with control or VEGF-C adenovirus. (A) Percentage and number of total CD4^+^ T cells. (B) Percentage and number of total CD8^+^ T cells. (C) Percentage and number of total activated CD44^+^ CD4^+^ T cells. (D) Percentage and number of activated CD44^+^ CD8^+^ T cells. (E) Percentage and number of IFNγ^+^ CD44^+^ CD4^+^ cells. (F) Percentage and number of IFNγ^+^ CD44^+^ CD8^+^ cells. (G) Relative expression of cytokines in the ear after control or VEGF-C adenovirus administration as detected by RT-PCR. Data are shown as mean ± SEM. Significance was determined using a two-way ANOVA paired with a Tukey’s multiple comparison test where *p<0.05 and **p<0.01.

